# Re-exploiting multiple RNA-seq data to identify transcript variations in *Podospora anserina*

**DOI:** 10.1101/2022.09.16.508226

**Authors:** Gaëlle Lelandais, Damien Remy, Fabienne Malagnac, Grognet Pierre

## Abstract

**Background:** Publicly available RNA-seq datasets are often underused although being helpful to improve functional annotation of eukaryotic genomes. This is especially true for filamentous fungi genomes which structure differs from most well annotated yeast genomes. *Podospora anserina* is a filamentous fungal model, which genome has been sequenced and annotated in 2008. Still, the current annotation lacks information about cis-regulatory elements, including promoters, transcription starting sites and terminators, which are instrumental to integrate epigenomic features into global gene regulation strategies.

**Results:** Here we took advantage of 37 RNA-seq experiments that were obtained in contrasted developmental and physiological conditions, to complete the functional annotation of *P. anserina* genome. Out of the 10,800 previously annotated genes, 5’UTR and 3’UTR were defined for 7,554, among which, 3,328 showed differential transcriptional signal starts and/or transcriptional end sites. In addition, alternative splicing events were detected for 2350 genes, mostly due alternative 3’splice site and 1,958 novel transcriptionally active regions (nTARs) in unannotated regions were identified.

**Conclusions:** Our study provides a comprehensive genome-wide functional annotation release of *P. anserina* genome, including chromatin features, cis-acting elements such as UTRs, alternative splicing events and transcription of non-coding regions. These new findings will likely improve our understanding of gene regulation strategies in compact genomes, such as those of filamentous fungi. Characterization of alternative transcripts and nTARs paves the way to the discovery of putative new genes, alternative peptides or regulatory non-coding RNAs.

## INTRODUCTION

If coding sequences define protein primary structures, messenger RNAs (mRNAs) direct their cytoplasmic expression. From pre-mRNA processing to translation initiation, their untranslated regions (UTRs) control most of the post-transcriptional gene regulation aspects, including nucleo-cytoplasmic transport, subcellular localization, mRNA stability and translation efficiency (1,2). To initiate gene expression at transcriptional start sites (TSS), transcriptional factors, histone chaperones (3) and chromatin remodelers (4) bind to cis-acting DNA sequences known as core-promoter, to recruit the RNA polymerase II complex. Conversely, transcription termination at specific transcription end sites (TES) prevent read-through transcription into adjacent genes, an acute concern in fungal compact genomes (5). Both 5’UTR and 3’UTR present a variety of canonical cis-acting elements that are bound by trans-acting elements (6,7). In addition, upstream ORF present in the 5’UTR are key regulators of translation (8). This combinatory repertoire tunes the composition of proteome (entire set of proteins) in accordance with developmental and/or metabolic needs of the cell. Several evidence suggests that the UTRs may harbor mutations that drives human traits and diseases (9), including cancer pathogenesis (10).

At a given core-promoter, the transcription may start from one of several TSS. Extensive studies performed on various human tissues established high-resolution transcription start sites maps (11). In animals, compilation of TSS localizations relative to gene expression identified two categories of core-promoters (reviewed in (12)). Core-promoters that show sharp initiation patterns, *i*.*e*. one main TSS, are found active in adult tissue-specific genes or terminally differentiated cell-specific genes, whereas core-promoters that show dispersed initiation patterns, *i*.*e*. multiple equally used TSS, are found active either for broadly expressed housekeeping genes or for developmental genes. In unicellular eukaryotes multiple or alternative TSS are often used to cope with changing environmental conditions. In the budding yeast, *in vivo* translation activities of alternative 5’UTR isoforms can vary by more than 100-fold (13).

In eukaryotes, chromatin accessibility is also a way to regulate gene expression. Condensed heterochromatin is less prone to transcription than opened euchromatin. To combine genome-scale functional information coming from both cis-acting elements (*i*.*e*. enhancers, promoters, TSS and TES) and histone modification patterns, schematic representations of model genes emerged for animals (14), plants (15) and some yeast species (16). Although a fairly large number of complete annotated fungal genome sequences is available (17), no such gene model has been built to date for filamentous fungi. Still, a recent assay for Transposase-Accessible Chromatin sequencing (ATAC-seq) performed in *Neurospora crassa* highlights the diversity of promoter structures and evidenced that histone acetylation and small RNA production are correlated with accessible chromatin, whereas some histone methylations are correlated with inaccessible chromatin (18).

Alternative splicing (AS) also regulates gene expression of eukaryotes. In animals, AS allows the generation of tissue- and time-specific isoforms, especially in brains. In *Drosophila*, the *Dscam* gene can generate over 38,000 distinct mRNA isoforms (19), which is more transcripts than the total number of genes in this organism (∼14,500). Notably, AS frequency is far less frequent in fungi than in animals, ranging from less than 1% in the budding yeast to 18% in the human pathogen *Cryptococcus neoformans* (20). Due to genomic features (few and short introns), intron retention (IR) is the most prevalent splicing type found in fungi (reviewed in (21)). However, studies performed in non-yeast fungi are limited.

*P. anserina* is a coprophilous ascomycete fungus that has been used as a model organism for almost a century (22). Its genome has been sequenced multiple times and watchfully annotated (23–25). However, no integrative genome-wide transcriptional landscape of *P. anserina* has been published yet. To do so, we took advantage of large and diversified sets of transcriptomic data and developed a customized annotation pipeline to map the 5’ and 3’UTRs genome-wide. We also evidenced the existence of alternative 5’and 3’ UTRs and described distinct types of alternative splicing events. Finally, novel transcriptionally active regions (nTARs) were searched and annotated, on which functional domain predictions were conducted to discover several putative new genes. We finally build a gene model that integrate the canonical *P. anserina* transcriptional features and the epigenomic landscape (26) in relation with gene expression status.

## RESULTS

### Collection of multiple RNA-seq data from various experimental conditions

A search for *P. anserina* in the SRA and BioProject repositories (27,28) returned 44 RNA-seq data from different studies on *P. anserina*’s life cycle (25), adaptation to carbon sources (29), response to bacteria (30) and senescence (31). Because it was generated by the SuperSAGE technology, this last dataset was excluded from the analyses. This left us with 37 RNA-seq from three studies, referred to as datasets A, B and C (Table 1). Altogether, these data cover a large variety of developmental states and growth conditions, which is important to increase the rate of transcriptionally active genes one might observe. Out of 1,054,787,963 reads in the 37 fastq files, 82.19% were mapped to the reference genome for which 10,800 CDS were annotated (23,24). Only 13 genes had no read mapped and 126 genes had only between 1 and 10 aligned reads (Fig. 1). Reads from dataset A alone, covering the entire life cycle, covered more than 99.7% of annotated CDS (respectively 31, 101 and 96 genes were not mapped in dataset A, B and C). This pool of dataset is then an interesting starting point to infer the transcript characteristics in *P. anserina*.

**Figure 1:**
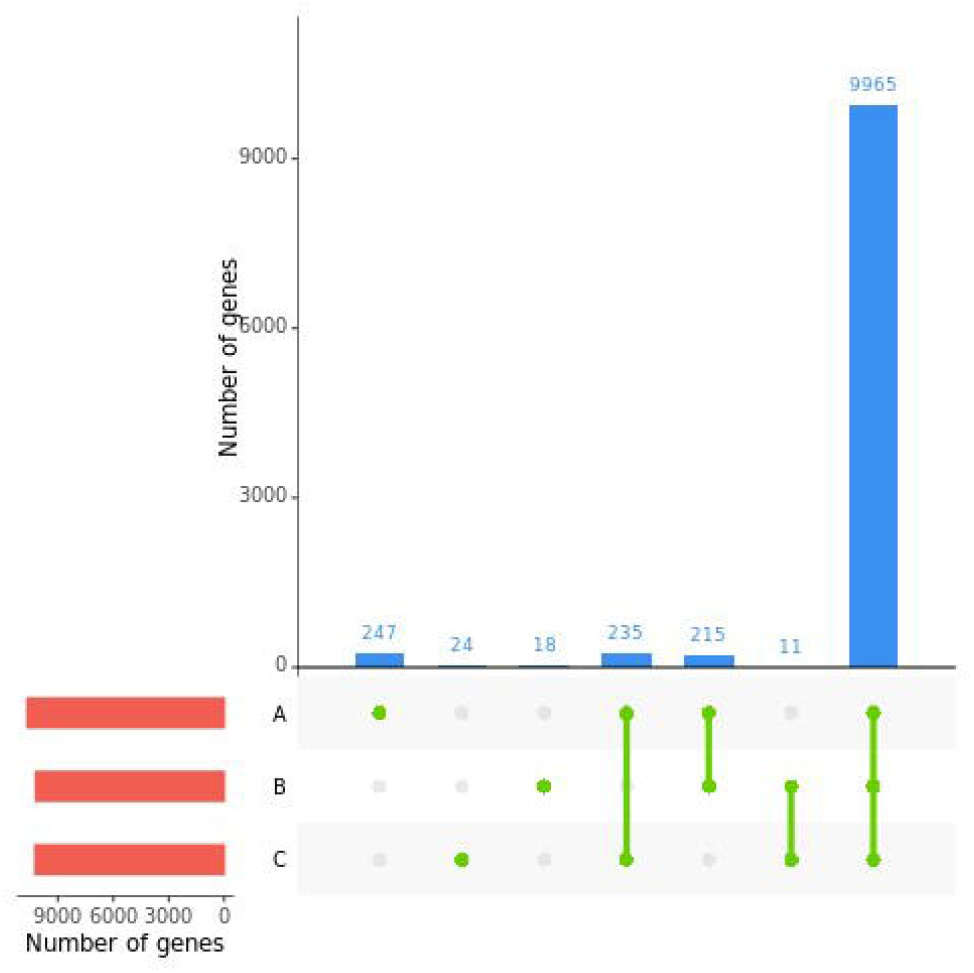
Number of CDS with mapped reads in the different datasets. Only CDS with at least 5 reads are shown here (n=10,724, 79 CDS are excluded). The red bar plot represents the number of genes with mapped reads in each dataset, the blue bar plot shows the number of genes with mapped reads shared or specific to one dataset as depicted with green points and lines bellow.

### Detection of TSS and TES for transcript related to already annotated CDS

Our first goal was to get a more accurate annotation of the *P. anserina*’s transcripts, related to the trustworthy annotated CDS in the genome. In this context, our rational was to consider that more accurate prediction for TSS and TES positions can be obtained with a high coverage of reads along transcripts. Therefore, the doubtfulness of the prediction decreases as the coverage increases (Fig. 2A). With that in mind, we developed a strategy in which we selected the most reliable transcript annotation, according to the read coverage (Fig. 2B, C). For a given gene, the multiple transcript annotations obtained from the 37 samples were sorted according to the average coverage value. Only those above a given threshold were next selected. The process was repeated for each gene to select the most accurate annotations. Hence, by using this Successive Coverage Values (SCV) method, only the most reliable annotations from all datasets were conserved.

**Figure 2:**
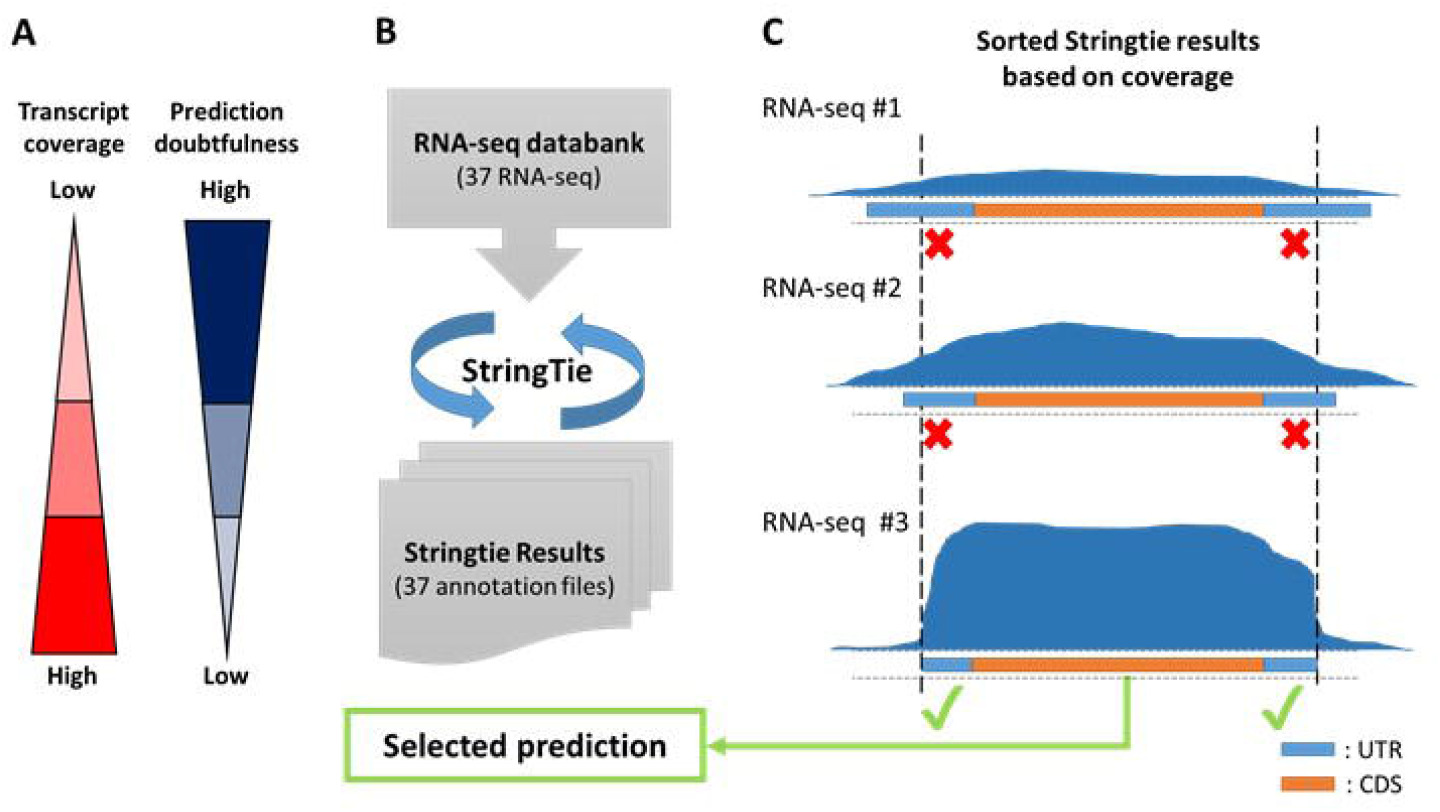
Schematic description of the Successive Coverage Values (SCV) methodology. **A**) This strategy is based on the assumption that for a given gene, the highest the coverage value, the less doubtful is the transcript prediction. **B)** All RNA-seq alignments are processed by the StringTie tool providing transcriptome annotation files in output. **C)** All transcriptome annotation files are compared gene by gene. First, transcript predictions are validated if they fully cover their associated CDS. Then, the average coverage from each validated prediction are compared and those above the most restrictive value are finally selected.

Applying this strategy on 10,803 predicted CDS of *P. anserina*, we could predict the transcript annotations for 7,554 genes (69.9% of the all set of CDS) (Table S1). The other CDS, for which no transcript prediction could be assigned, had very low coverage of reads. Thanks to the already available CDS annotations, we could get insight into the 5’ and 3’ UTRs characteristics. Of note, while 4,219 transcripts were predicted to have both a single TSS and TES, 3,335 genes got multiple transcript annotations (Fig. 3A-B). Most of the variations originated from both TSS and TES positions (Fig. 3C). Note that each dataset contributes significantly to the global annotation (Fig. 3D), highlighting the importance to work with a diversity of conditions, to get broadest transcriptional landscapes. We compare our annotation with the output of StringTie Merge and found our results more accurate as, in our case, StringTie Merge tends to fuse transcripts of closely located genes (FigS1).

**Figure 3:**
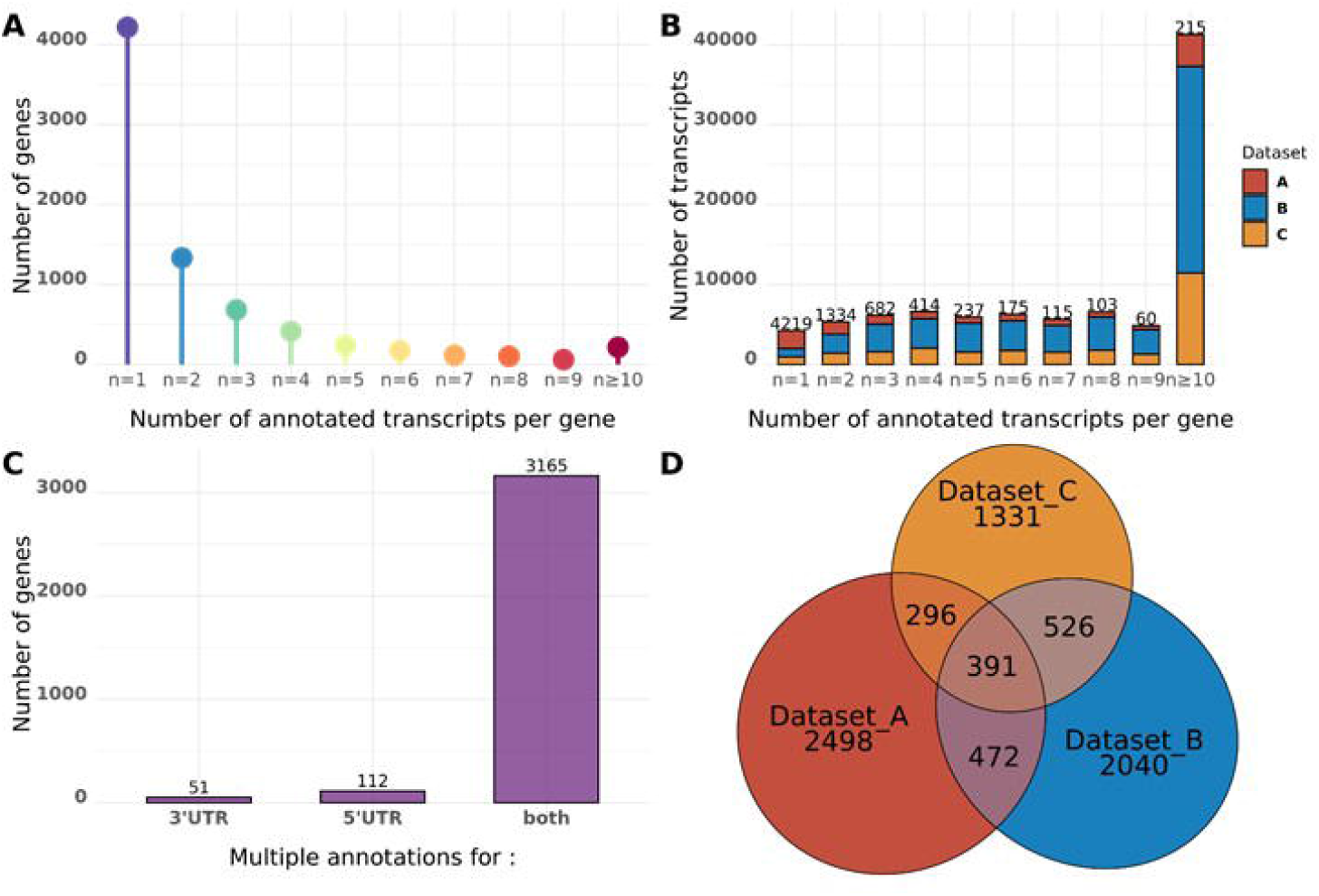
Transcript predictions of P. anserina’s genes. **A)** Distribution of genes with multiple transcript predictions. The number of genes in each category is written on top of the bar. Genes are divided in categories according to the number of predictions given by the SCV methodology. **B)** Distribution of the number of transcripts for each categories of genes. The number of genes in each category is written on top of the bar. **C)** Distribution of transcripts with different TSS and/or TES annotated for genes with multiple detected transcripts. D) Venn diagram of genes according to the dataset from which their transcripts are predicted. More than 75% of the genes with predicted transcripts have only one associated dataset (n = 5,869 genes).

The average sizes for the 5’ and 3’ UTRs were 275 bp and 303 bp respectively. When genes with single or multiple UTR are considered separately, the average size of UTRs do not extensively vary (Fig. 4A, Table 2). Indeed, we observed the most distant multiple TSS and TES are spaced with 156 bp and 114 bp in average respectively (Fig. 4B, Table 2), suggesting that if there are multiple transcription initiation or end sites, the transcripts do not display very different sizes. We also search for enriched sequence patterns located upstream of the defined TSS. Consistent with other fungal species, no clear TATA box was found.

**Figure 4:**
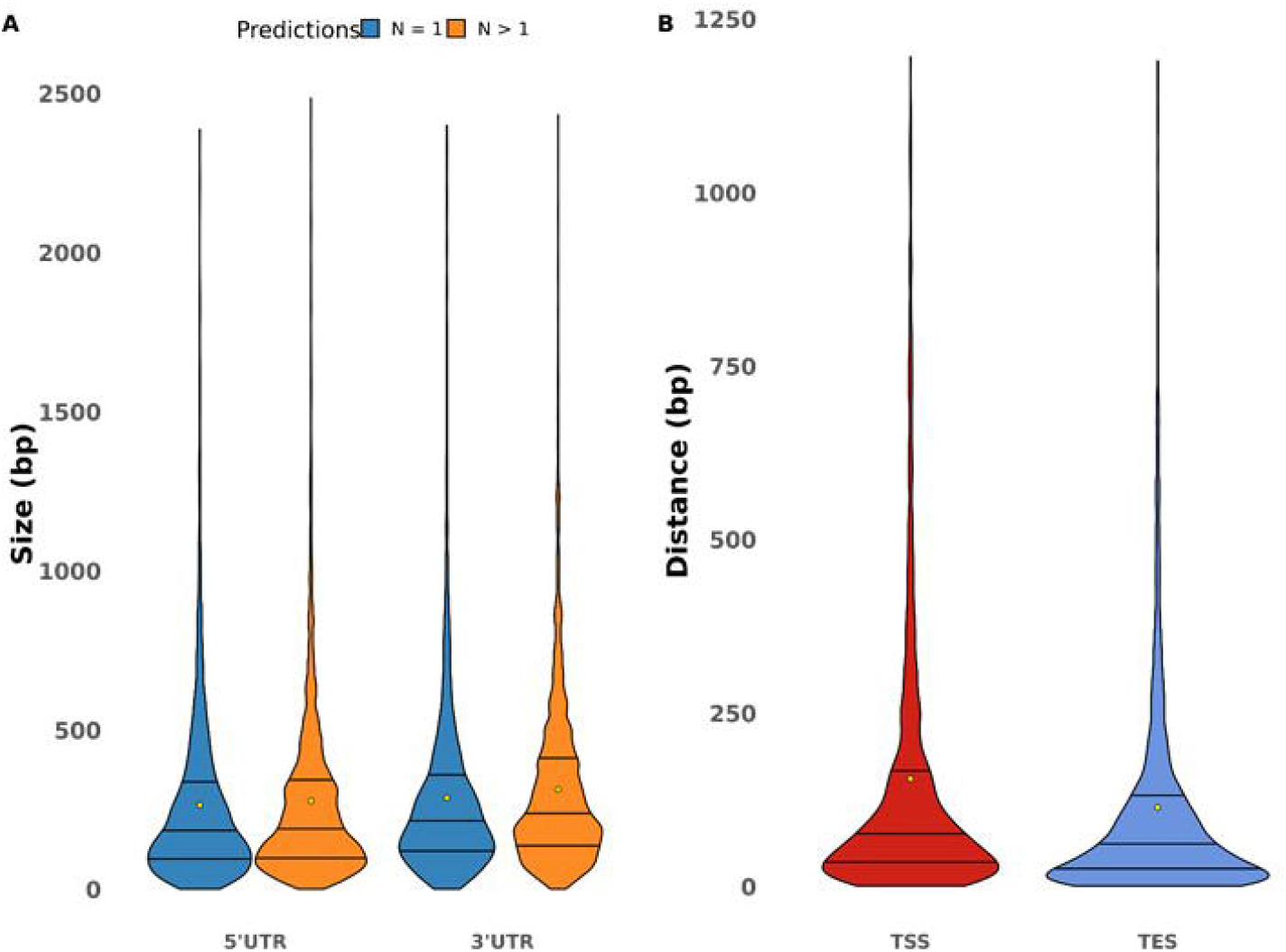
Size variation of UTRs. **A)** Summary of UTR sizes and violin plots representing the UTR size distribution. 5’ and 3’UTR size predictions are divided between genes with only one prediction. The yellow dots show the mean. For visualization purposes, the UTR bigger than 2 500 bp have been removed (n= 4 for unique 5’UTR, n=19 for multiple 5’UTR; n=2 for unique 3’UTR and n=19 for multiple 3’ UTR). **B)** Distance between the most distant TSS and TES. The mean distance values is marked with a yellow dot.

This allows us to describe the first average gene model in *P. anserina* shown Figure 5. The 5’ and 3’ UTR are 275 bp and 303 bp long, CDS is 1,483 bp long with an 80 bp long intron and the genes are spaced with 1,581 bp in average (Fig. 5).

**Figure 5:**
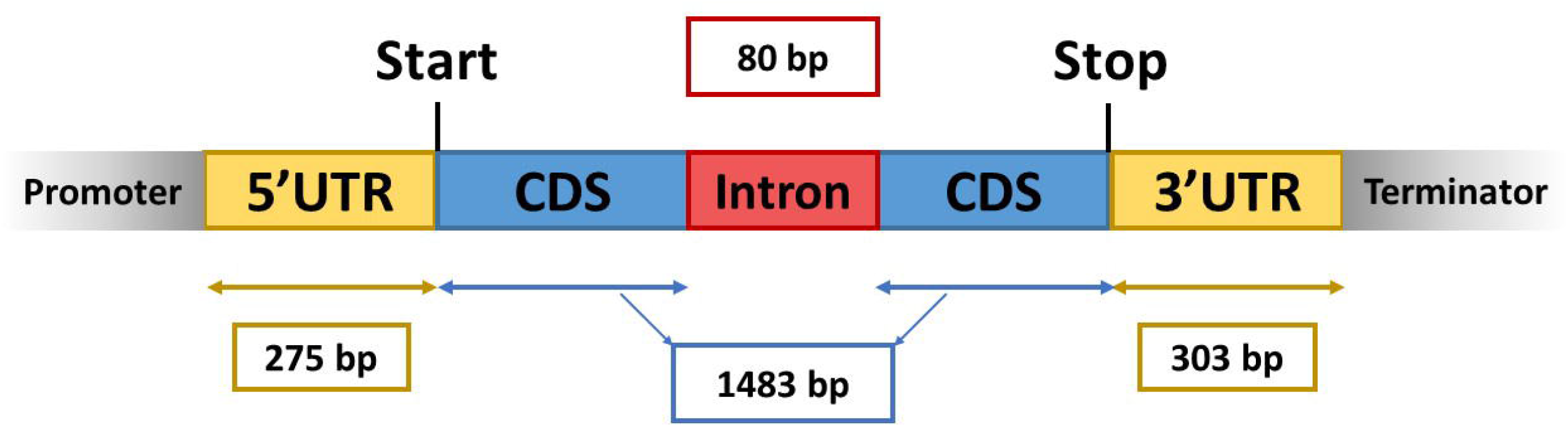
The average gene model of *P. anserina*. An average gene model was designed that includes all the new information generated here. TSS is expected at 275 bp upstream from the start codon and TES at 303 bp downstream from the stop codon. There are on average 1.49 introns per gene with a size of 80 bp. The mean CDS size is 1,483 bp. Yet, little is still known on promoter and terminator.

### Genome-wide schematic representation of average patterns of histone modifications in relation with transcription initiation

In order to validate our 5’UTR predictions, we took advantage of the ChIP-seq data that have been generated on histone marks in *P. anserina* (26). In mammals and plants, it has been established that H3K4me3 is enriched in the promoter region of active genes, whereas transcriptionally inactive gene promoters are rather marked with H3K27me3. We thus combined the enrichment of these two marks with our annotation for both transcriptionally active and inactive genes (Fig. 6). As a result, we could clearly observe that our predicted TSS positions fit with the peaks of H3K4me3 for expressed genes. Furthermore, the signal drop observed before the TSS, corresponds to the well described nucleosome free region (32). These observations support our predictions and show that data integration (RNA-seq and ChIP-seq) brings important information on gene organisation and epigenetic regulations of gene expressions.

**Figure 6:**
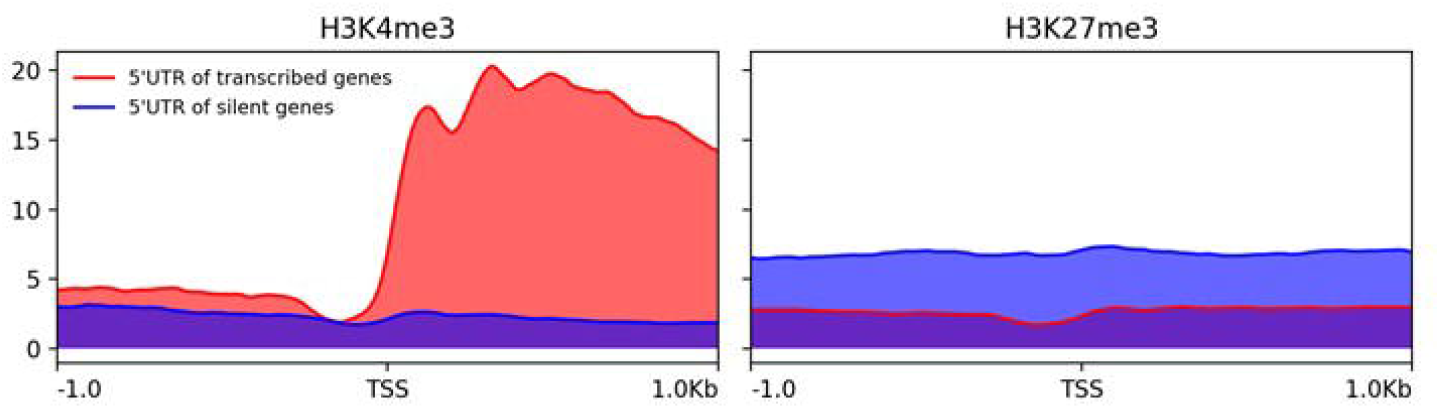
Integration of ChIP-seq data with the TSS annotations. **A)** H3K4me3 and **B)** H3K27m3 are mapped around the TSS of transcriptionally active and inactive genes. The x-axis represents a window span of 1 kbp upstream and downstream of the average TSS position. The y-axis represents the average value of the normalized number of reads mapped per bins (BPM normalization, bin size = 10).

### Detection of splicing sites and alternative splicing

In addition to the new annotations of UTRs, we used our RNA-seq dataset to validate the positions of introns in the current annotation of *P. anserina* genome. We could detect introns in the annotated UTRs for an important number of genes: 923 genes have at least one intron in their 5’UTR and 344 genes in the 3’UTR. Among them, 43 have introns in both UTRs. Furthermore, no information regarding the possible ASEs were available. We thus used the collected data to predict these ASEs. All four kinds of ASEs were detected: intron retention (IR), alternative 5’splice site (A5SS), alternative 3’splice site (A3SS), and exon skipping (ES) (Fig. 7) (TableS2). A total of 2350 genes were found subjected to at least one ASE. IR is the most frequent event; however, if the gene number is considered, A3SS represents the most frequent ASE detected in *P. anserina* with 1016 associated genes, followed by A5SS, IR and ES with respectively 758, 438 and 138 genes. A total of 278 genes could have isoforms with high combinatorial complexity (more than one ASE detected).

**Figure 7:**
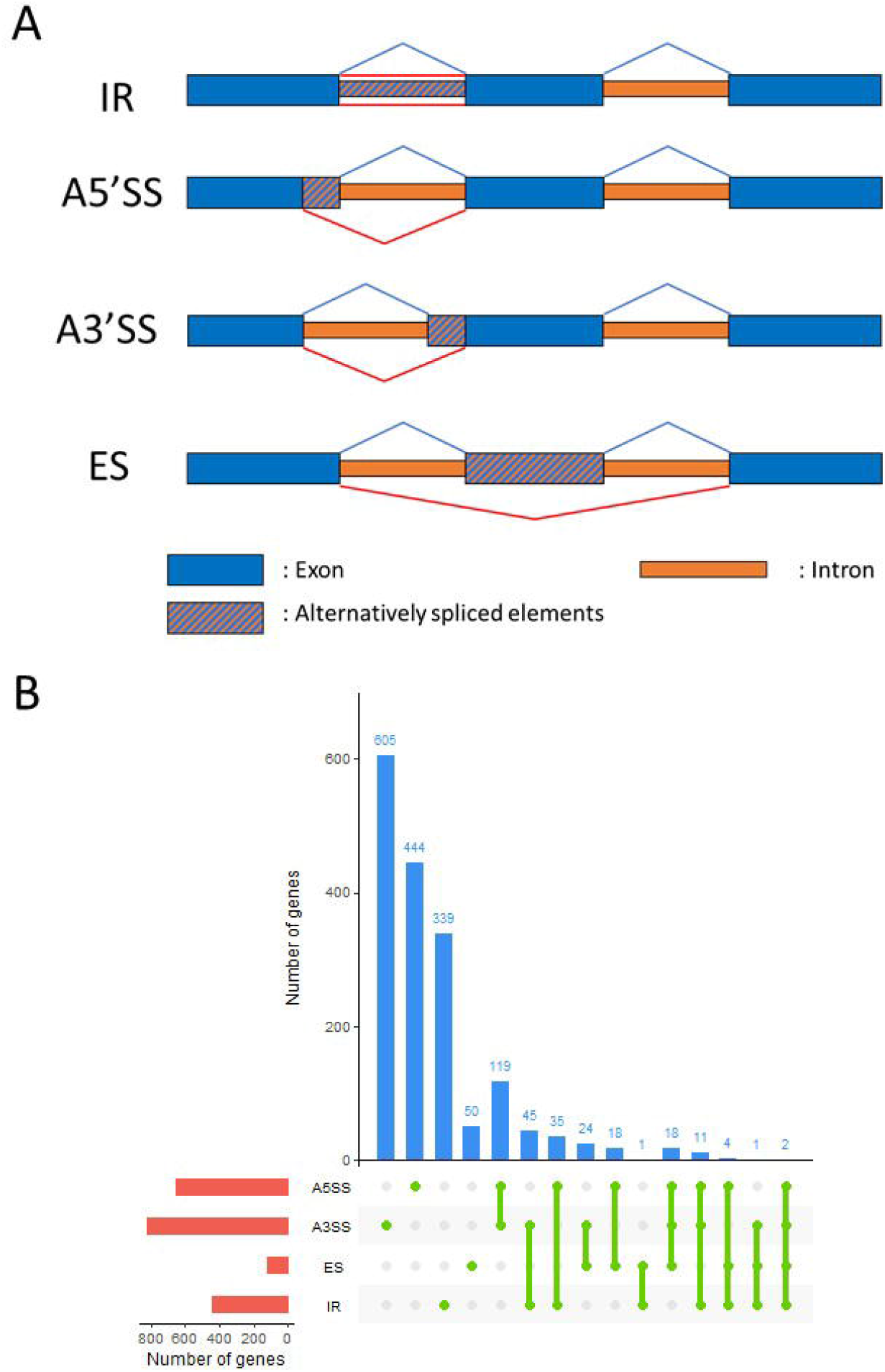
ASE detected in P. anserina coding transcripts. **A)** Representative example of four categories of ASE detected in *P*. anserina transcriptome. IR = intron retention, A5SS = alternative 5’ splicing site, A3SS = alternative 3’ splicing site, and ES = exon skipping (inspired from Kempken, 2013). **B)** Statistics of genes associated with ASE: The red bar plot represents the number of genes undergoing each type of ASE, the blue bar plot shows the number of genes undergoing the combination of ASEs depicted with green points and lines bellow.

### Identification of new transcripts, outside already annotated CDS

About 50% of reads mapped on the *P. anserina* genome were located in intergenic regions. They most likely correspond to novel transcriptionally active regions (named “nTARs” as in (33)). Therefore, we were able to detect 3,203 nTARs i.e. transcripts that do not fully cover already annotated gene. A significant part of them were very short (32% of nTARs shorter than 500 bp with a mean size of 1 kbp, while CDS length is app. 1.5 kbp long in average). Among all nTARs, 1,732 did not overlap any already annotated feature (Fig. S2) (with an average size of 1043 bp and 32% of them smaller than 500 bp). The 1,579 others were partially overlapping genes. Interestingly, we could detect introns in 55.8% of these 1,732 nTARs (N= 968) (Fig. 8) demonstrating production of processed transcripts by these potentially novel genes. Analysis of 332 nTARs longer than 1.5 kbp with the FGENESH gene prediction program yield 20 putative new protein-coding genes (Table S3). Domain prediction found 1 predicted gene with a putative rhodopsin C-terminal tail, transmembrane domains in 3 predicted genes and signal peptides in 2 predicted genes. One transcript was overlapping two sequences recently annotated as pseudogenes (25). No other protein domain was detected.

**Figure 8:**
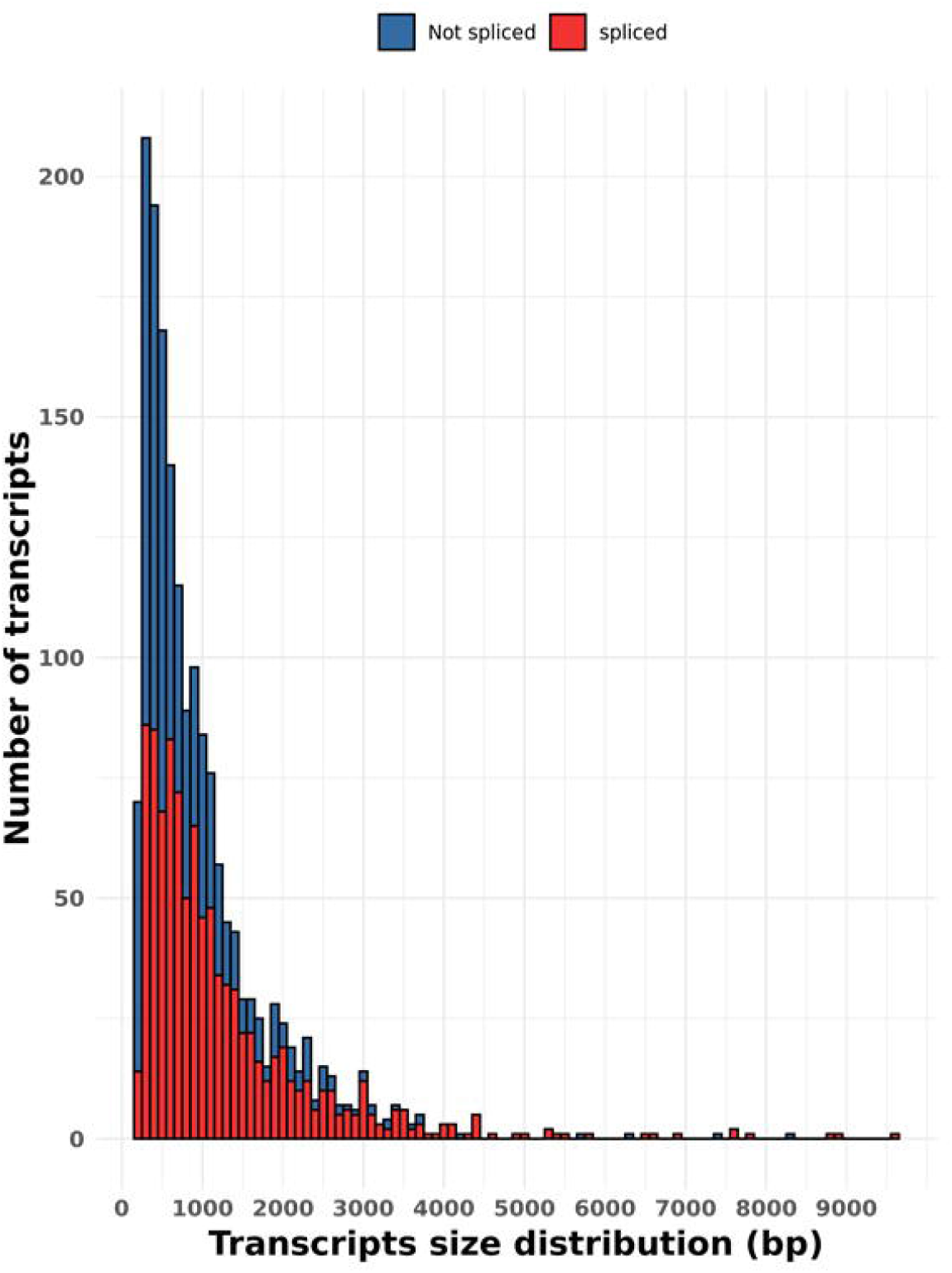
Transcripts predicted in non-coding regions. Size distribution of the 1,732 predicted transcripts in non-coding regions. The color shows the detection of splicing events in these transcripts.

One of the RNA-seq datasets was “stranded” (dataset B). This means that one knows from which strand the RNA molecule, which has been sequenced, originated. We thus used this dataset to seek nTARs overlapping previously annotated CDS but transcribed in the other direction, which we termed NATs (Noncoding Antisense Transcripts). NATs are long ncRNAs transcribed from the strand opposite to a protein-coding transcript, thus exhibiting sequence complementarity to mRNAs. We found 1472 NATs overlapping 452 genes (including 2 rRNA genes), 4 pseudogenes and 18 repeated sequences. Among these NATs on repeats, 7 were overlapping transposable elements which rules out a potential role of these NATs in silencing TEs, the other were found on segmental duplications.

## DISCUSSION

With this work, we completed the annotation of *P. anserina*’s genome by estimating transcripts size and variations using multiple RNA-seq data. Among the genes with mapped reads, we could make a trustworthy prediction of transcripts for more than two third of them, using a robust method, hence ensuring the reliability of the results. Although we detected multiple transcripts in 44% of the genes, we didn’t observed much variation in transcript size even with multiple TSS/TES. This is actually in line with previous observations. For example the *Masc1* gene in *Ascobolus immersus* has two TSS separated with 43 bp (35), whereas the *NiaD* gene in *Aspergillus nidulans* has two TSS separated with 72 bp (36). Usage of alternative TSSs in filamentous fungi has been described as transcriptional regulator in response of carbon source in *Aspergillus oryzae* (37) or translational regulator in regulating pathogenesis in *Metarhizium robertsii* (38). However, knowledge about how alternative TSSs affect gene expression is still nascent in filamentous fungi in contrast of what has been uncovered in mammals (39). In budding yeast (YeasTSS, (40)), a median of 26 transcript isoforms per gene were detected during regular growth conditions (41) and variable UTR sizes in different strains is linked with phenotypic variation (42). Usage of alternative TSSs and TESs is also involved in budding yeast cell fate transition. High resolution transcriptomic analysis evidenced elevated expression of alternative TSS and TES clusters in a stage-specific manner during yeast gametogenesis program and the mitotic cell cycle (43). Because, unlike yeasts, filamentous fungi present a syncytial organisation that cannot be synchronized, in-depth description of alternative TSSs and TESs remains challenging. However, our results show that over one third of *P. anserina* genes displays alternative TSS and/or TES usage. Moreover, when present, these alternative transcripts are specific of only one of the environmental conditions tested in this study. This suggests that the use of alternative TSS and/or TES also participates in *P. anserina* stage-specific gene expression and more generally to the resourceful ability of fungi for adaptation.

The genome average lengths of 5’UTR is quite similar across the diverse eukaryotic taxa, ranging from 100 to 200 bp with the size increasing during eukaryotes evolution (6), while genome average of 3’UTR lengths seems much more variable, ranging from 200 bp in plants and fungi to 800 bp in humans and 1000 bp in some vertebrates (44,45). Thereafter, analyses performed on larger sets of eukaryotic transcripts showed that although more variable than originally described, 5’UTRs average length is not as diverse as that of 3’UTRs in eukaryotic genomes (46). The *P. anserina* average size of both 5’ and 3’UTRs that we measured in this study were similar to those established for other fungal and non-fungal eukaryotes. In eukaryotes, the low size variability of UTRs contrasts with the very large size increase of intergenic regions during evolution. This intergenic space extension might be correlated with the necessity of a conserved “core promoter” structure, including the TATA box element. In *P. anserina* 5’UTRs we did not detect clear TATA box signature. This finding is consistent with previous observations showing that mots of fungal promoters do not contain a canonical TATA box (47). As a result, *P. anserina* likely uses the “scanning initiation” mode to start transcription rather than the “classic model” where most TSSs locate ∼30 bp downstream from the TATA box. Again, these different ways of initiating transcription might be correlated to the intergenic regions size, where a large sequence does not allow scanning and requires well defined sequences to recruit the polymerase. This new UTR annotation led us to search for UTR introns. As expected several UTRs were found to possess at least one intron although in a lesser extent than in human and plants (48,49). As important as the UTRs in gene expression are the introns present in these regions. Their splicing may affect both positively or negatively gene expression through various mechanisms such as mRNA export or nonsense-mediated decay (NMD). In eukaryotes, NMD degrades mRNAs containing premature stop codons as well as those containing an intron downstream of a stop codon, *i*.*e*. aberrantly spliced transcripts or 3’UTR intron-containing transcripts. Regulation of expression by mRNAs degradation is functional in *N. crassa* (Zhang and Sachs, 2015) and is expected to be functional as well in *P. anserina* since the NMD core components and the exon junction complex (EJC) are present (Table S4). The set of *P. anserina* genes for which we detected introns in their 3’UTR (∼3%) could therefore be prone to regulation by NMD.

We also looked at ASEs genome wide. In our prediction, the proportion of the different patterns of ASEs is in accordance with what has been observed in other filamentous fungi (50). Regarding the prevalence, we found almost 30% of the genes subjected to AS, which is far more than the 6% found in average for ascomycete fungi (20). However, this later estimation is based on ESTs and might underestimate the real number of ASEs (21). Indeed, recent RNA-seq showed ASEs in 24% of expressed genes in an oomycete (51) and 38% in a plant pathogenic ascomycete (52). One IR event was recently evidenced for the *PaKmt1* gene (26). This event can used as a positive control and has been indeed detected in our analysis supporting the robustness of our results. The physiological relevance of alternative splicing is still to be assessed in syncytial organisms but discovery of stage specific splicing events such as that of PaKmt1 suggests a finely regulated process in relation with developmental programs. In search for reliable ASEs we selected those present in at least two independent RNA-seq. Lifting this rules would allow us to detect stage specific ASEs but also expose to false positive.

In this study, we also identified thousands of novel transcripts. Some of them potentially encode functional proteins but the vast majority do not. Other comparable transcriptomic analyses expanded the annotated protein sets of *A. nidulans* and *U. maydis* by 2.9% and 2.5%, respectively (53,54). By comparison, the potential 29 new encoding proteins uncovered in this study represent only 0.3% of the previously annotated *P. anserina* CDS. This may be indicative of the good quality of its genome annotation. Among the non-coding nTARs, we detected potential antisense RNA that could also contribute to regulate gene expression. In fungi, non-coding RNAs, including natural antisense transcripts (NATs) are involved in development, metabolism, pathogenesis (55–57), etc. and can be expressed in a cell-specific manner (58). Some ncRNA/NAT are evolutionary conserved among related smut fungi, which suggests conservation of the corresponding ncRNA/NAT functions (53). In *N. crassa* and *A. nidulans*, antisense transcripts represents ∼5% and ∼14% of the annotated protein-coding loci, respectively (54,59). Since only one of the 37 RNA-seq is strand-specific and therefore suitable for antisense transcripts, it is too preliminary to quantify the importance of ncRNA/NAT contribution to gene expression. However, this study revealed the first evidence of expression anticorrelation between asRNAs and downstream CDSs.

By collecting information on UTRs and alternative splice sites, as well as identifying novel protein-coding genes and new isoforms, this study, among others, contributes to a better understanding of the molecular basis that governs gene expression in fungi. By overlaying these integrated transcriptomic data with our previous epigenomic data, we were even able to propose the first filamentous fungus gene model, as to what already exists in animals and plants.

## METHODS

### Collection, alignment of RNA-seq data and transcriptome annotation

RNA-Seq data were downloaded from the Sequence Read Archive (SRA) (28) at the accession number ERR2224046 to ERR2224051 (25), SRR3197700 to SRR3197711 (60), SRR6960207 to SRR6960225 (29). Each fastq file was mapped onto the genome of *Podospora anserina* S *mat*+ (23–25) using HISAT2 version 2.1.0 (61) with default parameters. In order to make sure that all the data are of equivalent quality (e.g. no RNA extremity degradation in one dataset) raw read coverage was checked on constitutively expressed genes (Fig S3) and verified to be consistent across studies at least for the beginning and end of the transcripts. Output alignments files were respectively processed by the StringTie program version 1.3.5 (62) with parameters:–g 5 –c 10 (--rf only for RNA-seq from dataset B) and default values for the remaining parameters.

### Read quantifications

Reads counts were performed for each alignment files using htseqcount version 0.6.0 with the following parameters: --stranded no (RNA-seq A and C) --stranded reverse (RNA-seq B) --mode intersection-non-empty.

### Transcript annotation

The annotation files were processed by the Successive Coverage Values (SCV) method using custom-made scripts. The SCV algorithm selects transcript predictions that fully overlap only one annotated CDS by their genomic coordinates. Then, respectively for each genes, transcript predictions from different RNA-seq are compared by their average coverage value to a discrete scale of threshold (from 10 to 20000 reads). Transcript predictions above the most restrictive threshold are selected to annotate a new single gene model. In order to keep valuable information without redundancy for genes with several transcripts predictions, the longest UTRs are selected and all alternative start and end signal of transcription are annotated considering their RNA-seq.

### Integration of ChIP-seq data

ChIP-seq data normalized coverage value were from (26). Active genes and inactive genes were selected respectively as the 800 most expressed and 800 less expressed in the M4 condition from dataset A as calculated in (25). This sample was chosen because it is the closet from the conditions used for the ChIP experiments.

### Detection of new transcripts

All StringTie transcript predictions with an average coverage equals or higher to 10 that did not intersect any coding or non-coding element of the current annotation were annotated as novel non-coding transcripts. Then, nested transcripts were merged using bedtools version 2.29.2 (63) with default parameters. Spliced non-coding transcripts were retrieved by intersection of TopHat junctions (TopHat2 version 2.1.1, Trapnell et al., 2009) with genomic coordinates of non-coding transcripts.

Gene prediction was made with FGENESH (65) and domain prediction with InterProScan version 5.52-86.0 (66)

### Detection of motif in promoters region

The 200 bp sequences upstream of each TSSs have been extracted using bedtools’ flank and getfasta tools (63). Motif search has been performed using MEME with default parameters (67).

### Alternative splicing events detection

To obtain information of alternative splicing events that occur in *P. anserina*, we used TopHat2 version 2.1.1 (64) with all default parameters but --min-intron-length 30, --max-multihits 5 and specifically –segment-length 21 for dataset A and --library-type fr-firstrand for dataset B. Exon-exon junctions annotations were then processed to filter out low-confidence exon-exon junctions (independent RNA-seq ≥ 2 and coverage ≥ 5 in at least one RNA-seq for A5SS, A3SS and ES).

For IR, we quantified aligned reads on CDS and introns annotations following the method above (see Read quantifications). Introns annotations were segmented by 8 bp bins to assess coverage variability. Then, retained introns were selected applying four thresholds (T1, T2, T3 and T4): 1) average coverage per bin higher than T1, 2) standard deviation of coverage per bin lower than T2, 3) overall expression of the associated CDS higher than T3 and finally 4) ratio between average coverage per bin of the intron and the overall expression of the associated CDS higher than T4. Different association between threshold values were tested, to finally retain: T1=30, T2=20, T3=200 and T4=0.1. These values allowed to properly select a positive control, i.e. the intron Pa_6_990.G_intron_1 which was expected to be retained in the experiments ERR2224048 and ERR2224049 (26).

### Detection of the NATs

Potential NAT (Noncoding Antisense Transcripts) were extracted from the StringTie outputs, obtained with dataset B (see section “Collection, alignment of RNA-seq data and transcriptome annotation”). They are transcripts which 1) have coordinates that overlap (entirely or partially) only one annotated CDS in Podospora anserina genome and 2) are found on the opposite strand.

## LIST OF ABBREVIATIONS

mRNA: Messenger RNA;
UTR: Untranslated region;
TSS: Transcription start site;
TES: Transcription end site;
CDS: Coding sequence;
IR: intron retention;
A5’SS: Alternative 5’splicing site;
A’3SS: Alternative 3’ splicing site;
ES: Exon skipping;
ASE: Alternative splicing event;
nTARs: Novel transcriptionally active regions;
SCV: Successive coverage values

## DECLARATIONS

## Ethics approval and consent to participate

Not applicable

## Consent for publication

Not applicable

## Availability of data and materials

All scripts and annotation files can be found at: https://github.com/Podospora-anserina/transcript_annotation_2022

## Authors’ contributions

Design analyses: GL, FM, PG. Analyze data: DR, GL, PG. Wrote the paper: FM, PG.

## Acknowledgements

We thank Robert Debuchy for fruitful discussions and precious experimental help. We thank Daniel Gautheret for his suggestions and critical reading of the manuscript. We are also thankful to Cecile Fairhead, Philippe Silar and Stephane Le Crom for their expertise and fruitful discussions.

## TABLE AND FIGURES LEGENDS

**Table 1: Composition of the RNA-seq databank**. 37 RNA-seq in total parted in 3 datasets were collected from public database (SRA and BioProject, accession number provided). The *Cs* strain genome from dataset C only differs from the strain *S* at the *het-s* VI locus. Overall, the 37 RNA-seq used in the analysis represent 19 unique experimental conditions. The heterogeneity of this pool of dataset provides better chance to observe genes in their active expression state.

**Table 2: Summary of UTRs characteristics. A)** Summary of UTRs sizes for both unique and multiple TSS/TES prediction When multiple UTRs, data are calculated from all UTRs from all genes. **B)** Summary of distance between most distant TSS and TES for each gene with multiple 5’UTR and/or 3’UTR. Mean, median and maximum sizes are expressed in base pairs

**Table S1: Transcripts predicted in all the RNA-seq** dataset **by StringTie and selected with the SCV method**.

**Table S2: List of alternative splicing event detected for each gene**.

**Table S3: Gene prediction from the putative new transcripts**

**Table S4: Genes coding components of NMD and EJC**.

**Fig S1:**
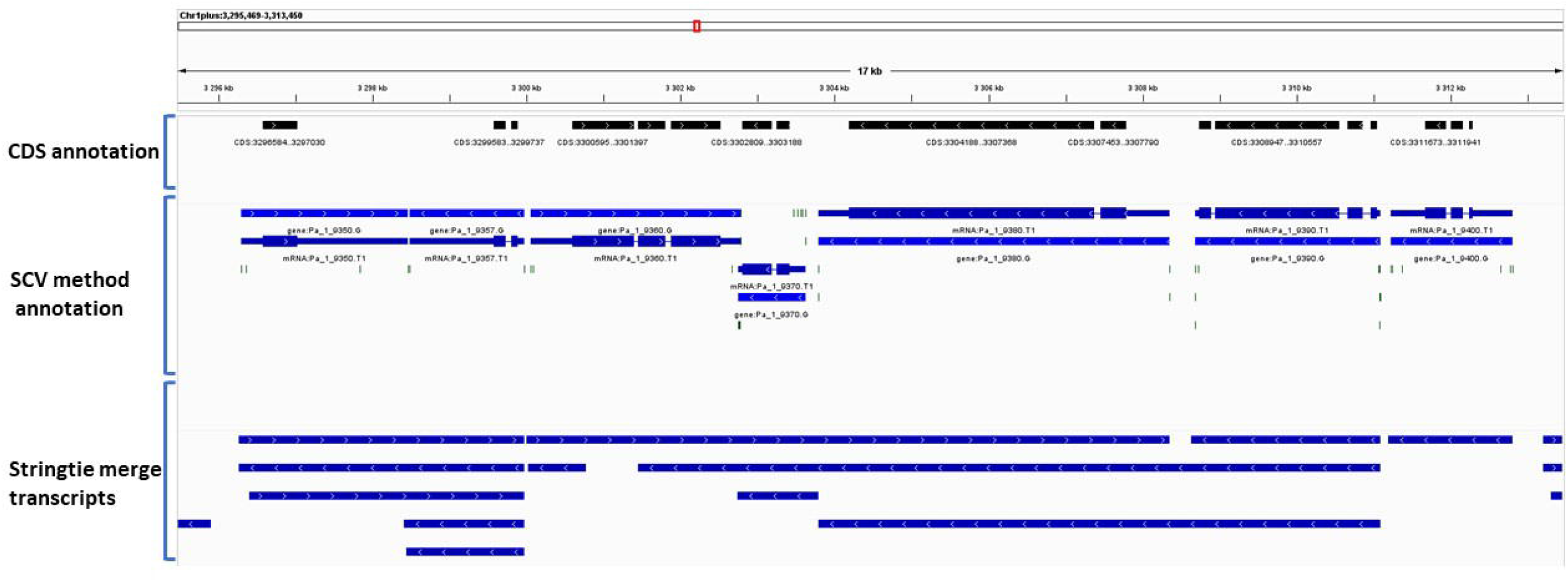
Comparison of the SCV method annotation and StringTie merge results. The different annotation files have been visualize with IGV. This screenshot is representative of the rest of the genome. The small horizontal black lines show the positions of the TSS and TES for the SCV method annotation.

**Fig S2:**
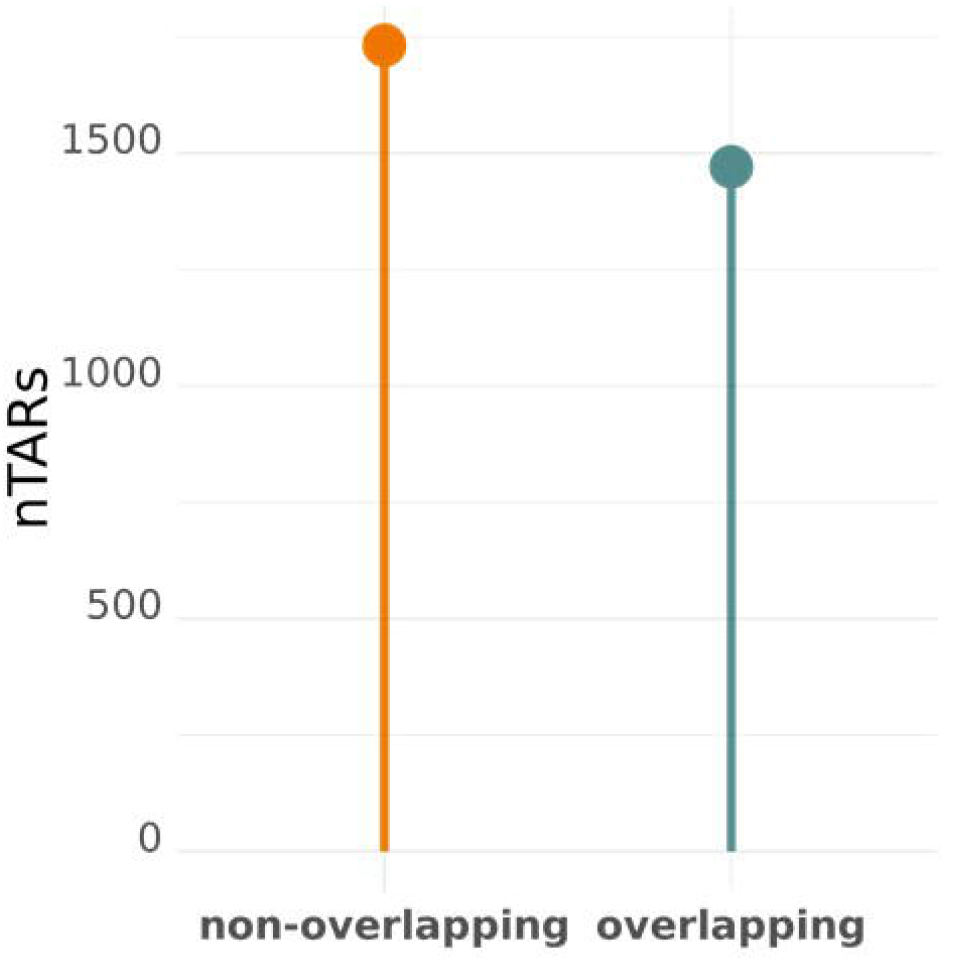
Quantification of nTARs overlapping or not previously annotated features.

**Fig S3. RNA-seq quality control**. The read coverage of each sample is shown for four constitutively transcribed genes. The CDS and gene annotation is shown on the bottom of each plot. No major discrepancy between the dataset were observed.

